# Impact of C-terminal amino acid composition on protein expression in bacteria

**DOI:** 10.1101/751305

**Authors:** Marc Weber, Raul Burgos, Eva Yus, Jae-Seong Yang, Maria Lluch-Senar, Luis Serrano

**Author notes:** Correspondence (L.S.).

## Abstract

The C-terminal sequence of a protein is involved in processes such as efficiency of translation termination and protein degradation. However, the general relationship between features of this C-terminal sequence and levels of protein expression remains unknown. Here, we identified C-terminal amino acid biases that are ubiquitous across the bacterial taxonomy (1582 genomes). We showed that the frequency is higher for positively charged amino acids (lysine, arginine) while hydrophobic amino acids and threonine are lower. In highly abundant proteins, the C-terminal residue is more conserved. We then studied the impact of C-terminal composition on protein levels in a library of *M. pneumoniae* mutants, covering all possible combinations of the two last codons. We found that charged and polar residues, in particular lysine, led to higher expression, while hydrophobic and aromatic residues led to lower expression, with a difference in protein levels up to 4-fold. Our results demonstrate that the identity of the last amino acids has a strong influence on protein expression levels and is under selective pressure in highly expressed proteins.

## Introduction

Protein sequence is shaped by many evolutionary constraints, acting at different levels of the gene expression process. Identifying which sequence features determine the efficiency and accuracy of protein expression has been the subject of intense research. Sequence variations previously believed to be neutral, such as the choice of synonymous codons, have been shown to be under selection (1), revealing new mechanisms interacting with the translation process. Most studies have focused on the region close to the N-terminal, where some of the most important mechanisms of translation initiation occur (2–4).

Much less is known, however, about the potential evolutionary pressures acting at the C-terminal, apart from basic protein function and structure. Early studies showed a differential preference for specific codons upstream of the stop codon in *Escherichia coli* (5–8) and *Bacillus subtilis* (9, 10) proteins. The properties of the last two amino acids were shown to modulate the efficiency of translation termination at the UGA stop codon context in *E. coli* (7), in particular for highly expressed genes. Also, several C-terminal sequence motifs were found to induce stalling of translation termination (11, 12). Besides, degradation signals have been identified at the C-terminal of proteins in a variety of bacterial species (13), such as the ssrA tag which targets proteins to the ClpXP protease in *E. coli* (14). Thus, changes in the efficiency of translation termination and recognition of the C-terminal region by the protein degradation machinery are two potential mechanisms that could drive preferences in the C-terminal composition of proteins.

Translation is one of the most energy-intensive processes in the cell, consuming about 40% of the cellular energetic resources in fast-growing bacteria (15). Thus, sequence features at the C-terminal that lead to variations in protein abundance, by modulating either translation or degradation rates, are likely to be under selection. However, the impact of C-terminal composition on protein abundance remains largely unknown. Many studies have identified sequence features that influence translation efficiency, but most of them have focused on the 5’ end region or the bulk of the coding sequence (4, 16, 17). In these studies, the use of synthetic libraries of a reporter gene with randomized sequence has proved as a robust approach to evaluate the functional impact of sequence variation. A similar approach applied to variations in the C-terminal region would provide useful information to identify sequence features associated with higher or lower protein abundance.

Here, we investigated the impact of C-terminal sequence composition on protein expression. Firstly, we leveraged the considerable increase in the past decade in the number of available bacterial genome sequences (18) to study C-terminal composition biases of 1582 genomes across the bacterial taxonomy. Such a large-scale comparative analysis allowed us to reveal the universality of the sequence compositional patterns, as well as preferences which seem to be species specific, and unveil their association with protein function and protein abundance. Secondly, we experimentally assessed, in a high-throughput manner, the influence of C-terminal composition on protein expression levels in the model organism *Mycoplasma pneumoniae* using the ELM-seq technique (19). We built a random library of the *dam* reporter gene with varying C-terminal sequence, covering all possible combinations of the last two codons and the six nucleotides following the stop codon. By measuring the expression levels of all variants, we showed that the identity of the last two amino acids have a strong impact on protein abundance. We validated these results by varying the last residue of a different protein in the same species. Overall, our results show that in bacteria, the C-terminal residue of protein sequences modulates protein expression levels and is under selective pressure in highly expressed proteins.

## Results

### Analyzing C-terminal compositional biases in bacteria

#### C-terminal amino acid and codon composition in bacteria is biased

We investigated biases in codon and amino acid composition of the C-terminal region of bacterial protein sequences. We retrieved all protein sequences from the RefSeq database (20), using the reference and representative genome collections, in order to achieve a broad coverage of bacterial species across taxonomy. To avoid over-representation of duplicated proteins within the same bacterial species, we removed proteins which presented both a high overall sequence identity and a high identity of their C-terminal region (see Methods). We obtained a database of approximately 4.8M protein sequences covering 1582 genomes and 1516 species, which we used as a starting point for all the following analyses.

When studying all species in the bacterial kingdom, we found that the amino acid composition at the last position upstream of the stop codon differed significantly from the bulk amino acid composition (Fig. 1A). In particular, positively charged amino acids were enriched at position −1, with the frequency of lysine and arginine being 2.32 times (Fisher two-sided exact test p ≈ 0 within numerical error) and 1.76 times (p < 7.7e-30) higher, respectively. In contrast, the occurrence of threonine was 2.25 times (p < 2.2e-308), and that of methionine, 2.02 times (p < 7.1e-51) lower. Due to the large number of sequences considered in this analysis, all biases were significant with extremely small p-values, even after correcting for multiple tests. Interestingly, we observed a gradient in the intensity of the biases for all amino acids that showed a statistical difference at the C-terminal, except for threonine that was specifically depleted at the −1 position. This gradient was more evident in the case of arginine and lysine, whose biases decreased from a maximum value at the C-terminal position towards the bulk, with an odds ratio still significant for lysine of 1.16 (p < 2.3e-23) at position −20. All hydrophobic amino acids, except phenylalanine, were found to be disfavored at the last position, with fold changes in frequency ranging from 0.49 to 0.87 times (p < 7.5e-6). The amino acid frequencies detected at positions −1 and −2 differed from those found in disordered regions in proteins, indicating that the preferences observed are not due to the C-terminal being in general unstructured in proteins (□^2^ test p ≈ 0, Fig. S2).

**Fig. 1.**
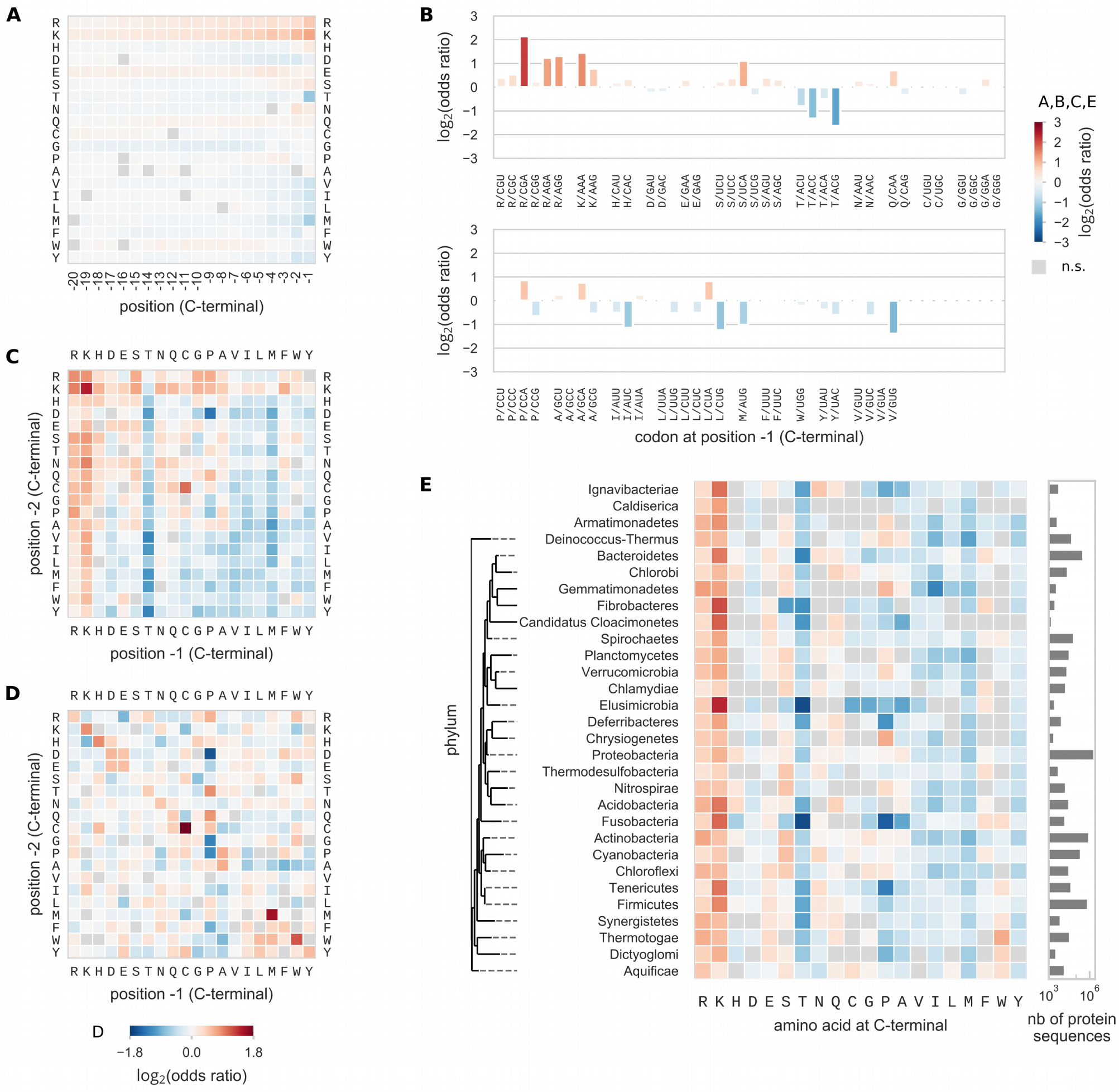
Biases in C-terminal protein sequence composition in the bacterial kingdom. Amino acid and codon composition at the C-terminal of bacterial protein sequences shows higher (red) or lower (blue) frequency when compared to their frequency in the bulk of sequences (same color code for panels A, B, C, and E). Significance of the biases were tested using exact Fisher test and multiple-tests correction with 5% false discovery rate within each plot category. (A) Position-specific amino acid composition biases for the last 20 amino acids at C-terminal. (B) Codon composition bias at the last position. (C) Composition bias of the last two amino acids compared to the frequency of the amino acid pair in the bulk. (D) Epistasis between the last two amino acids. Frequency of the pair was compared to the expected frequency if the two positions were independent. In this case significance was tested with the binomial test and the same multiple tests correction. Color code has a smaller range than in other panels. (E) Amino acid composition bias at the position −1 for individual phyla. Phyla were ordered following an approximate phylogenetic tree.

To explore possible cooperative effects we compared the frequency of an amino acid pair at the last two positions to its expected frequency under the assumption that both positions are independent (null model) (Fig. 1C). The deviation from the expected frequency, or epistasis, revealed that many of the pairs of repeated residues were more frequent than expected; in particular Cys-Cys-Stop (odds ratio 5.16), Met-Met-Stop (odds ratio 2.93), Trp-Trp-Stop (odds ratio 2.24), His-His-Stop (odds ratio 1.84), and Lys-Lys-Stop (odds ratio 1.80), (binomial test, p ≈ 0 for all). These five amino acid pairs also exhibited a positive epistasis in the bulk (Fig. S3) (odds ratio 1.29-1.53), suggesting that part of the observed positive interaction was not specific to the C-terminal. In addition, because Cys, Met, Trp and His were also the least frequent amino acids in general, a small number of proteins with a conserved functional motif that include those dipeptides could easily lead to the overrepresentation of the pair. In particular, the Cys-Cys pair presented the strongest positive epistasis effect. In the family of metal sensor proteins, binding to metal ions is often mediated by multiple cysteine thiolates (21, 22). While the metalloregulatory protein families are diverse in structure and functions, some of them possess a conserved cysteine-rich motif close to the C-terminal (23). Indeed, at least 396 out of the 2034 proteins in our database that possess a CC-Stop motif belonged to orthogroups related to metalloprotein families. On the opposite, the amino acid pairs that exhibited the most negative epistasis were some of the Xaa-Pro-Stop combinations Asp-Pro-Stop (odds ratio 0.33), Gly-Pro-Stop (odds ratio 0.43), Pro-Pro-Stop (odds ratio 0.44), Phe-Pro-Stop (odds ratio 0.54) (binomial test, p ≈ 0 for all). Two of these dipeptides (Asp-Pro and Pro-Pro) were previously shown to induce the strongest level of translation stalling and tagging by the ssrA ribosome rescue system in *E. coli* (11), providing a possible explanation for this negative selection.

In order to explore if composition biases are conserved across the bacterial phylogeny, we grouped bacterial genomes into taxonomic clades at the level of phyla, and analyzed composition biases of sequences from each reduced set of bacterial species (Fig. 1D). Overall, the main biases at the C-terminal position were present in virtually all phyla. Interestingly, proline was found to be enriched in 12 phyla and depleted in other 16 phyla (e.g. 0.23 odds ratio in Tenericutes, 0.17 in Fusobacteria). The same analysis at a finer level of the taxonomy (Fig. S6, Fig. S7) showed that the biases for proline varied greatly across taxonomic clades. Interestingly, the biases across phyla for threonine anti-correlated with the biases for lysine (pearson *r* = −0.84).

#### Pattern of C-terminal codon biases and its relationship to the stop codon context

We then asked whether those biases were restricted to specific codons or were rather independent of the identity of the synonymous codon (Fig. 1B). We found that both codons encoding for lysine (AAA and AAG) were enriched at the −1 position at fairly similar levels (odds ratios 3.13 and 2.43). Similarly, all four codons coding for threonine were depleted, despite some variations in their odds ratios. Interestingly, only three (CGA, AGA and AGG) out of the six arginine codons were strongly enriched, with the highest odds ratio for the CGA codon (4.44). In the group of hydrophobic amino acids that were found to be depleted (valine, isoleucine, leucine, methionine, tyrosine and tryptophan), some synonymous codons were more strongly underrepresented, although in general the majority of these codons were disfavored (11 out of 17). We emphasize that these preferences were not related to differences in synonymous codon usage, since all biases were computed with respect to the frequency of codons in the bulk. Interestingly enough, with the exception of arginine AGA and AGG codons and lysine codons, in all other amino acids there was a strong preference for an A at the third position.

We reasoned that some of these variations in C-terminal codon biases could be related to specific interactions with the stop codon. Thus, we analyzed the biases in C-terminal codon composition in each of the three stop codon contexts, both at the level of bacterial kingdom (Fig. 2) and at the level of individual phyla (Fig. S4). Overall, codons that presented strong enrichment (depletion) were enriched (depleted) in all stop codon contexts. However, some variations in codon biases were also observed. We found that the identity of the last base of the C-terminal codon had an influence on codon biases depending on the stop codon context (Fig. S10). In particular, NNA codons were more often enriched than other codons in all three stop codon contexts (distribution of the log2(odds ratio), independent t-test p < 4e-24). This preference was exacerbated in the UGA context, where NNA codon were clearly favored over NNG codons (p < 1.8e-49). In this respect we should mention that ending with an A in the context of a UGA stop codon creates an overlapping starting AUG codon, while the other bases will results in the less favoured UUG, GUG, CUG start codons.

**Fig. 2.**
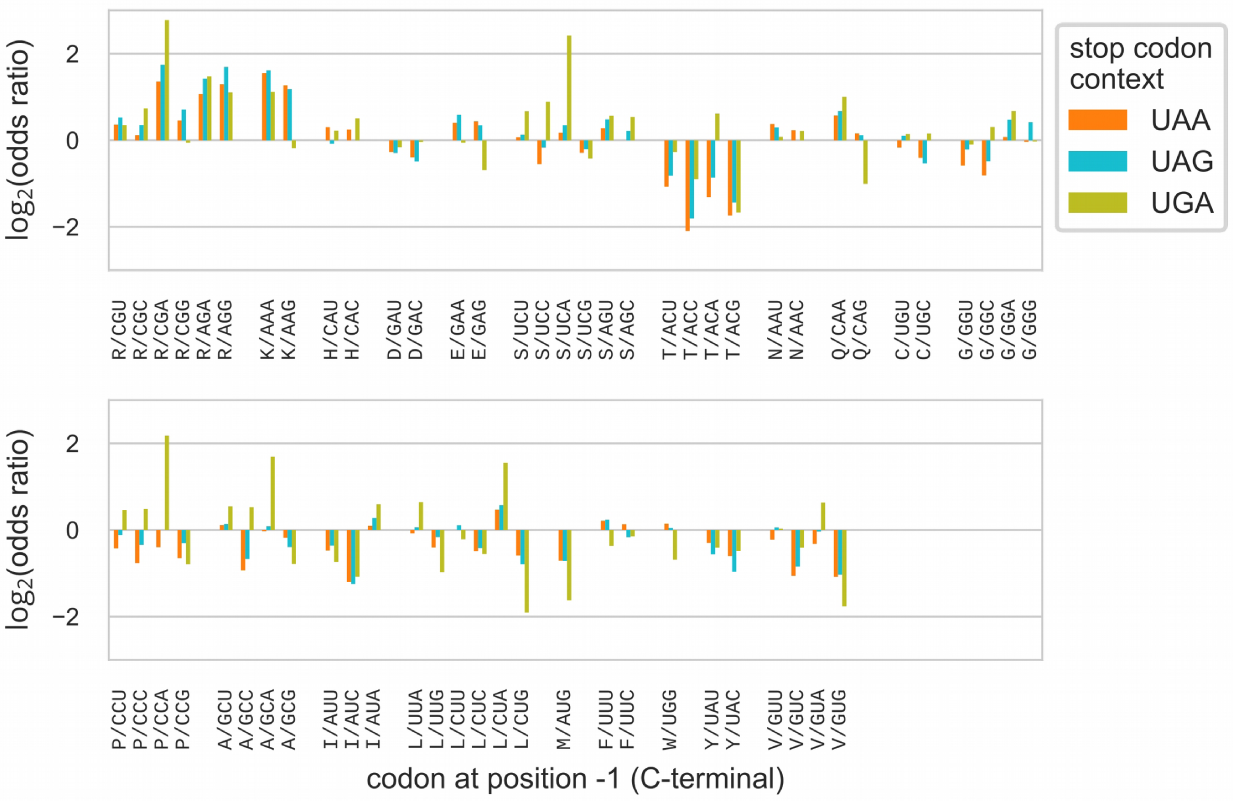
Codon composition biases at C-terminal and stop codon context. In the bacterial kingdom, protein sequences are first classified by their stop codon context. Codon frequency at C-terminal is then compared to the bulk codon frequency within each stop codon context class. Significance of the biases were tested using exact Fisher test and multiple-tests correction with 5% false discovery rate within each class.

#### Pattern of C-terminal amino acid biases is qualitatively independent of functional category

Next, we asked whether the biases in C-terminal amino acid composition are specific to some protein functional classes. We hypothesized that some of the biases could be driven by a small group of proteins with amino acid composition different from the average. In order to explore this hypothesis, we analyzed protein functional groups in two ways.

First, we studied whether the enrichment for lysine and arginine at the C-terminal region could be due specifically to transmembrane proteins. It has been suggested that the orientation of transmembrane domains is determined by the enrichment of positively charged residues in cytoplasmic loops, rather than in periplasmic loops, a mechanism known as the positive-inside rule (24, 25). Thus, transmembrane proteins whose C-terminal domain is cytoplasmic might present an enrichment of lysine and arginine compared to their frequency in the bulk of the sequence. Indeed, positive charges have been shown to be enriched in the C-terminal cytosolic region of transmembrane proteins in *E. coli* (26). We classified proteins as membrane or cytoplasmic based on computationally predicted localization for a selection of 364 bacterial species, and computed the C-terminal amino acid composition biases for each of the two classes (Fig. S12). Positively charged residues were found to be strongly enriched in the last 10 positions of the C-terminal of membrane proteins (mean odds ratio K, 2.10, R, 1.69). The same biases were weaker for cytoplasmic proteins (mean odds ratio K, 1.57, R, 1.22). Hydrophobic amino acids were found to be depleted in both protein categories, although more strongly in membrane proteins (mean odds ratio for A, V, I, L, M, F, W, Y, 0.72 for membrane, 0.84 for cytoplasmic). Apart from these differences, we found a similar pattern of biases at position −1 for membrane and cytoplasmic proteins (Fig. S12C), including depletion of threonine, methionine and hydrophobic residues, and enrichment of lysine and arginine. Thus, while membrane proteins have a higher frequency of positively charged residues at the C-terminal, they only partially contribute to the global amino acid composition biases observed at the level of all proteins.

Second, we systematically classified proteins into functional categories by assigning each protein sequence to a Cluster of Orthologous Groups (COG) category. We computed the composition biases within each of the 23 functional categories, by comparing the frequency of amino acids at the C-terminal to the bulk frequency of sequences in the same category (Fig. S13). The previously observed general biases were maintained in the vast majority of the functional categories. Importantly, these biases were maintained despite differences in the bulk frequency of some amino acids between categories. For example, ribosomal proteins contain many positively charged residues that are essential for their interaction with RNA (27), and as a consequence, proteins in the J category “Translation, ribosomal structure and biogenesis” have a higher frequency of lysine in the bulk (6.01% compared to 4.23% in average for the other categories). However, at the last position of the C-terminal, lysine is found at even higher frequency (16.2%) than in the bulk, resulting in a strong enrichment (odds ratio 3.02).

Therefore, the main amino acid biases observed at the last position of the C-terminal are qualitatively independent of functional category.

#### C-terminal amino acid identity is associated to protein abundance

One possibility is that the observed C-terminal biases could be driven by an underlying mechanism affecting protein abundance, such as translation termination efficiency or protein degradation. If this is the case, we would expect to observe differential biases for proteins that are highly abundant than for lowly abundant proteins. We examined the association between protein abundance and C-terminal biases in 13 bacterial species for which at least 40% of the proteins had experimental abundance value from the PaxDB database (28). We categorized proteins into low (percentiles 0-20), medium (percentiles 20-80) and high (percentiles 80-100) abundance, and computed the amino acid composition biases in each group (Fig. 3). Highly abundant proteins showed a stronger enrichment of Lys but not for Arg, and depletion of Thr, Pro and Cys at the C-terminal (position −1) compared to lowly abundant proteins. In addition, amino acid biases at position −3, with the exception of Cys, were very similar between the two abundance categories. Thus, the identity of the C-terminal amino acid at position −1 could be in part related to protein abundance.

**Fig. 3.**
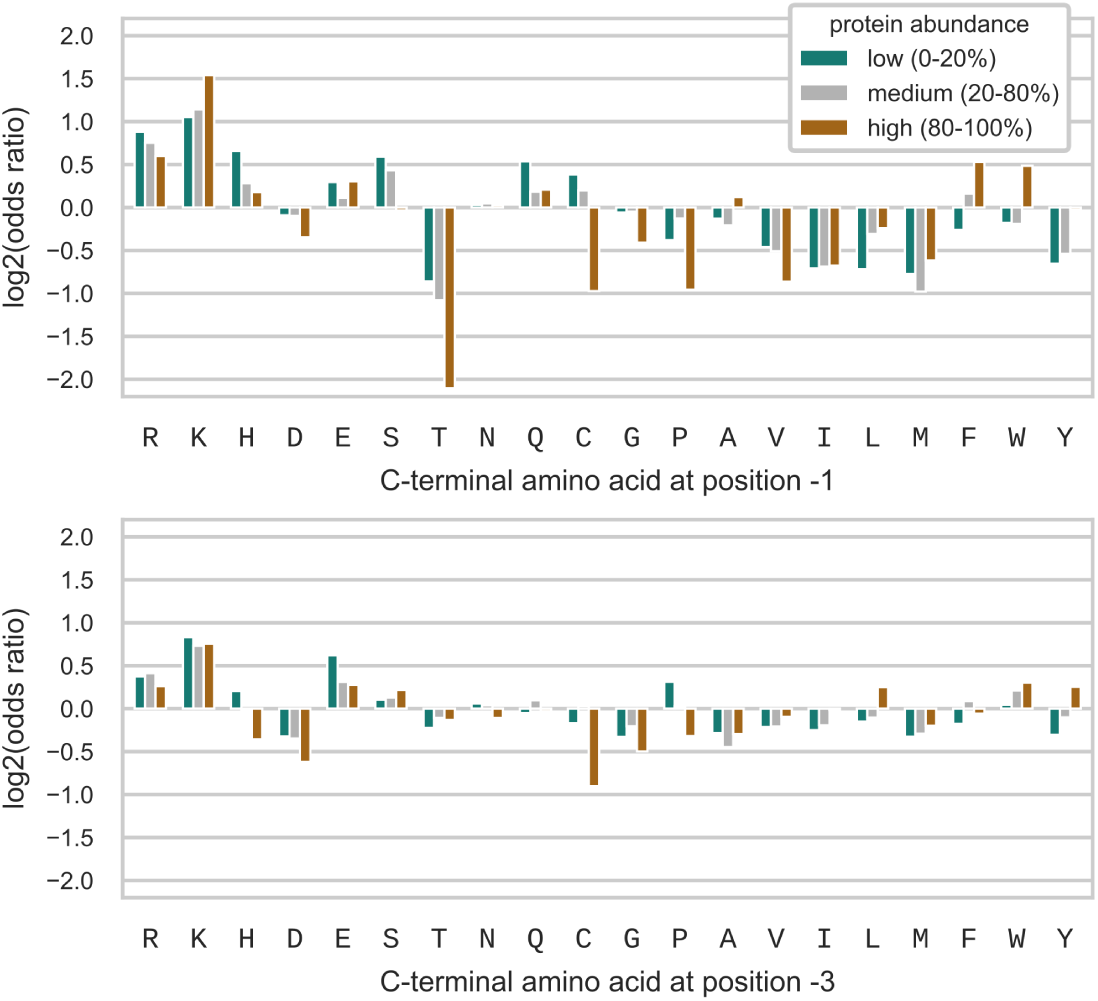
Protein abundance and C-terminal amino acid identity. Proteins were categorized into low (percentiles 0-20, green), medium (percentiles 20-80, grey) and high (percentiles 80-100, brown) abundance, and C-terminal amino acid composition biases, with respect to the bulk frequency of each category, at position −1 (upper plot) and position −3 (lower plot), were analyzed for each of the three categories.

#### Last residue is preferentially more conserved within variable C-terminal regions of orthologous groups

We then investigated whether the composition of the C-terminal residues is under selective pressure. Proteins are exposed to multiple evolutionary constraints that act on their sequence and may potentially overlap with and mask the selective pressure on the C-terminal residues. In order to unveil such a position-specific selection, we selected orthologous groups (OGs) that presented a non-conserved and variable in length C-terminal region (see Methods and Fig. 4). In these cases, we assumed that residues in the C-terminal region were not essential for protein structure or protein function, and hypothesized that any position-specific conservation pattern would be the consequence of such a selective pressure.

**Fig. 4.**
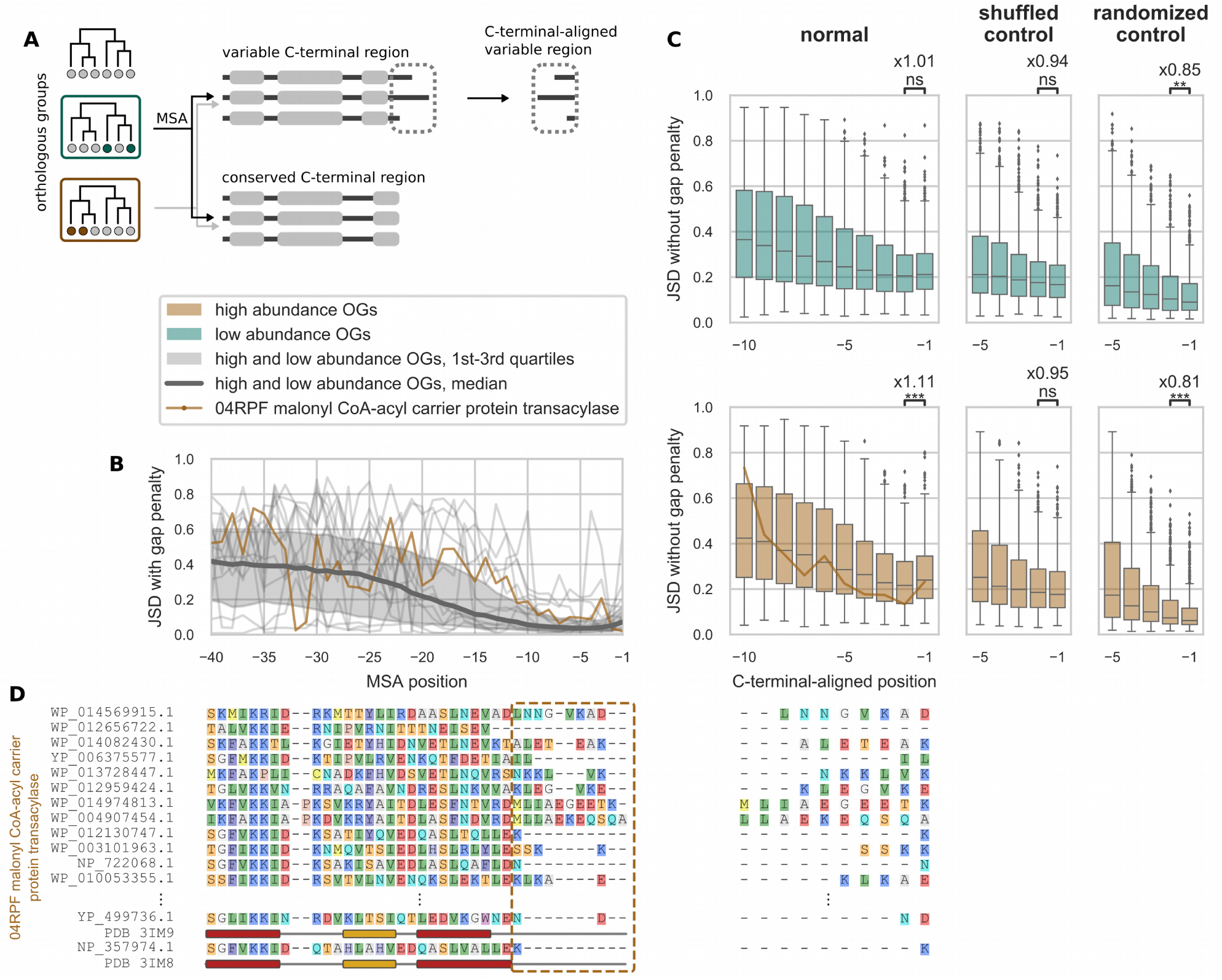
Conservation profile of orthologous groups with variable C-terminal region. (A) Orthologous groups were first classified into high or low abundance classes based on partial protein abundance data. Orthogroups (OG) with variable C-terminal region were selected, and their variable region was then aligned to the right. (B) Profile of Jensen-Shannon divergence (JSD) conservation score with gap penalty of the selected OGs. (C) For each OG, variable region sequences were aligned to the C-terminal and the JSD conservation score without gap penalty was computed. Distributions of the conservation score for each C-terminal-aligned position for the low (upper plots) and high (lower plots) abundance OGs showed a significant increase between the −2 and −1 position in the high abundance class (W.M.W. test p < 3.8e-04), while no significant change was found for the low abundance class (W.M.W. test p < 0.23). Shuffled sequences showed no significant change between the two positions. Random sequences of the same length exhibited a decrease, due to the increase in the number of non-gap residues in the last column. (D) Example: Multiple Sequence Alignment (MSA) of malonyl CoA-acyl carrier protein transacylase (MCAT) OG exhibits a variable C-terminal region in the last 11 columns. Only 14 sequences are shown out of the 68 non-redundant sequences that were selected to compute the MSA. Residues were colored following the RasMol amino acid color scheme. Secondary structure annotations indicate alpha helix (red) and beta strand (gold).

For each orthogroup, we aligned sequences within the variable region to the C-terminal (right-aligned), and computed a conservation score for each column (see Methods). In the case of the low abundance OGs (Fig. 4C), we observed that the distribution of the conservation score for position −1 was not significantly different from the distribution at position −2 (W.M.W. test p < 0.23, fold change of mean 1.01). For OGs in the high abundance class, however, the distribution of the conservation score at position −1 was significantly higher compared to the position −2 (W.M.W. test p < 3.8e-04, fold change of mean 1.11). In the absence of any selective pressure in the variable region (null hypothesis), one would expect the opposite trend, i.e. a decrease in conservation. We validated these results by performing two randomized controls (shuffled and randomized sequences) as described in the Methods section.

Thus, these observations support the existence of a selective pressure, not related to protein function or structure, specifically acting on the identity of the C-terminal residue of highly abundant proteins.

### C-terminal amino acid identity impacts protein expression levels in *M. pneumoniae*

As described above, the identity of the C-terminal residues could be related to protein abundance and be under selection in highly abundant proteins, suggesting that it could play in role in determining protein expression levels. To see if this could be the case we selected a genome reduced organism with a simplified degradation (40) and translation (39) machinery, *M. pneumoniae*, to remove as many confounding factors as possible. In particular, it has only one release factor (RF1) recognizing the UAA and UAG stop codons (29), with the UGA codon coding for tryptophan. In addition, it lacks a peptidoglycan sacculus and no carboxypeptidase has been identified in this species (30). We designed an experimental assay to measure, in a high-throughput manner, the expression levels of the *dam* reporter gene (DNA adenine methylase) with a variable C-terminal region, using the ELM-seq method (19) (see Fig. 5A and Methods). Expression levels, as measured by this method, are approximately linear to the protein abundance (19), which facilitates quantitative interpretation of the results. We built a random library of the *dam* reporter gene where six random nucleotides were added at the C-terminal of the Dam wild-type sequence, such that all combinations of the last two codons were covered. Six random nucleotides were also added downstream of the UAA stop codon, in order to study their influence, if any. Transcription of the *dam* gene was driven by either a strong or a weak promoter, resulting in two different average levels of expression.

**Fig. 5.**
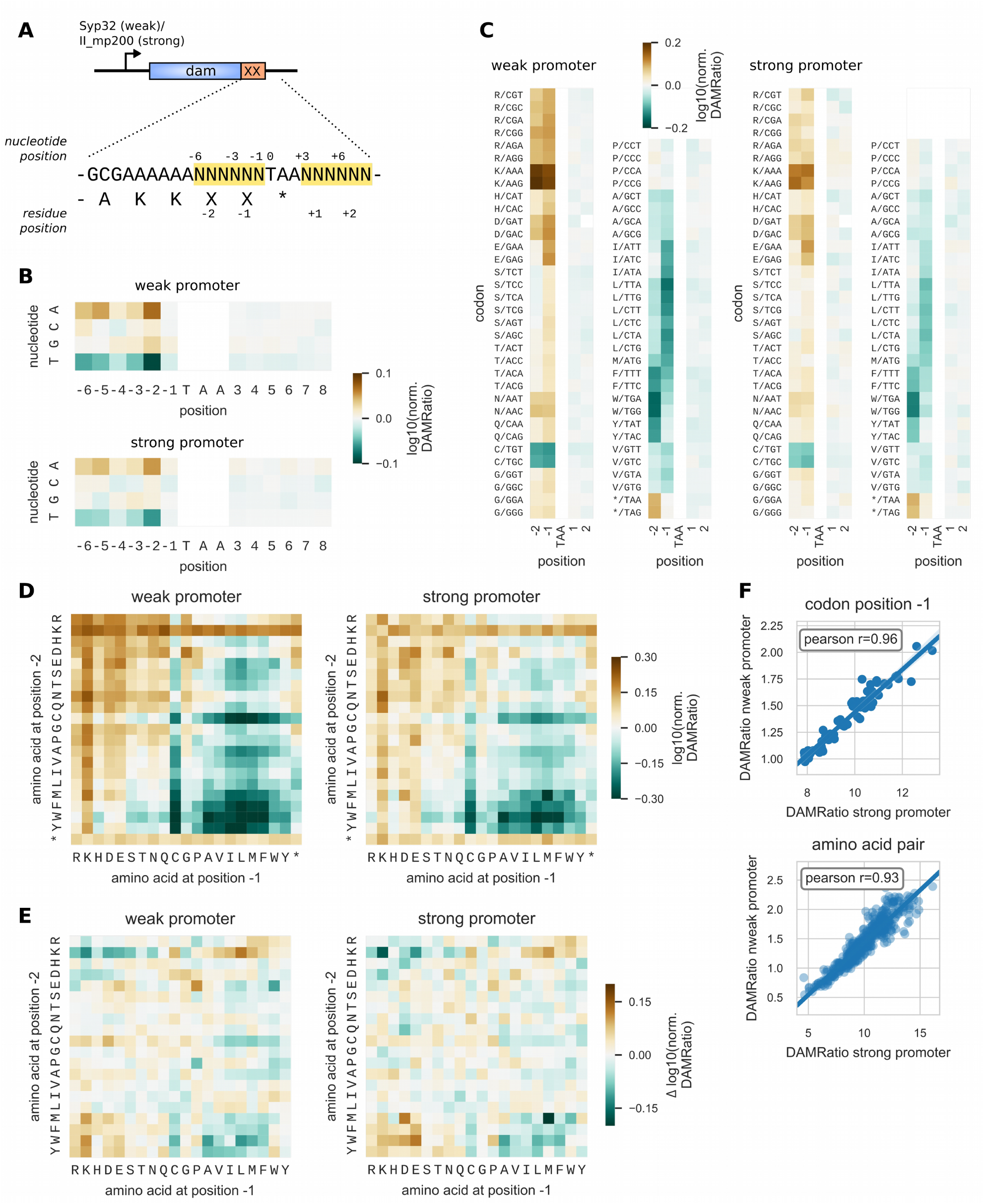
Protein quantification of a randomized C-terminal library using ELM-seq. (A) A C-terminal library was created with the *dam* reporter gene, where the last 6 nucleotides before and the 6 nucleotides after the stop codon were randomized (indicated by Ns). Two versions of the library were created using either a weak or a strong promoter. Expression levels in *Mycoplasma pneumoniae* were measured by ELM-seq, a technique that combines DNA methylase as a reporter with methylation-sensitive restriction enzyme digestion and high-throughput sequencing. Protein expression level readout is reported as the log10 of the DAMratio relative to the average. (B) Effect of nucleotide identity at each position on expression levels. (C) Effect of codon identity at in-frame codon positions −2, −1 (before the stop codon), +1 and +2 (after the stop codon) on expression levels. (D) Effect of C-terminal amino acid pair identity (last two codon positions before the stop codon) on expression levels. (E) Cooperativity in the amino acid pair effect on expression levels (epistasis), as measured by the difference between expected and measured log10(DAMRatio). (F) Correlation of expression levels between weak and strong promoter libraries, at the level of codon at position −1 and at the level of amino acid pair.

We first examined the results in the weak promoter library. We analyzed the expression levels (DAMRatio) at the level of individual nucleotide identity for each of the random positions (Fig. 5B). The nucleotide at the first two positions of each codon had the strongest influence, while the 3rd nucleotide position had little influence, which suggests a similar expression level among synonymous codons. As expected, the nucleotide composition downstream of the stop codon had no influence on the expression levels.

We examined how expression levels varied depending on the codon identity at in-frame positions −2 and −1 (last two codons) (Fig. 5C). Clearly, amino acid identity at both positions −1 and −2 had a strong influence on expression levels. Positively charged amino acids (Lys, Arg) at either position led to a higher expression. In particular, the presence of one lysine residue at one of the two positions increased expression level by 1.48 to 1.55 fold compared to the average. Hydrophobic amino acids Ala, Ile, Leu, Met, Val, Phe, Trp and Tyr at either position produced in general lower expression levels (for the two positions, mean normalized DAMRatio 0.86, standard deviation 0.08). Cysteine also led to lower expression levels (normalized DAMRatio 0.76-0.79). Unexpectedly, we did not observe any significant changes in expression in the presence of proline or threonine. The random library also contained the stop codons UAA and UAG at positions −2 and −1, which acted as an internal control. In the presence of a stop codon at position −1, the expression level was identical to the average (x1.01), recapitulating the effect of all possible C-terminal codons at position −2. When a stop codon was located at position −2, we observed an expression higher than the average (x1.23), due to the two lysine codons at position −4 and −3 of the wild-type sequence.

We quantified the cooperativity (epistasis) between the amino acid identities at the two positions (Fig. 5E) by computing the ratio *Q* between the measured expression level of the pair to the expected expression level assuming that the effect of the two positions are independent. Most of the pairs showed almost no cooperative effect. Epistasis for the pairs KK-Stop, RK-Stop and KR-Stop was negative, which most probably resulted from the saturation of the cumulative effect of C-terminal amino acids on protein expression levels. In fact, the presence of one lysine at either position −2 or −1 resulted in higher than average expression levels regardless of the identity of the other residue (Fig. 5D). Negative epistasis was also found for the pairs [M,F,W,Y]-[A,V,I,L,M,F]-Stop (mean *Q*, 0.88, standard deviation 0.06), which led to the lowest expression levels (Fig. 5D). The pairs PP-Stop and RP-Stop exhibited negative epistasis (*Q =* 0.96, norm. DAMRatio 0.86), while DP-Stop and EP-Stop showed positive epistasis (*Q* = 1.10, norm. DAMRatio 1.18). Apart from the lower expression for the PP dipeptide (norm. DAMRatio 0.81), these results disagreed with the depletion of DP and EP observed in the genomic analysis, and the fact that these two motifs were shown to induce high level of ssrA tagging in *E. coli* (11). Overall, the expression levels varied up to 4.42-fold in the weak promoter library (3.51-fold in the strong promoter library), between the pair with the lowest expression, WL-Stop (MM-Stop), and the pair with the highest expression KQ-Stop (NK-Stop).

The results were highly reproducible, as shown by the strong correlation of the normalized DAMRatio between the weak and the strong promoter libraries (Fig. 5E), both at the level of the codon identity at position -1 (pearson correlation 0.96), and at the level of amino acid pair (pearson correlation 0.93). In the strong promoter library, the pattern of changes in expression was similar but less pronounced than in the weak promoter library, probably due to saturation effects.

We examined whether other sequence factors could play a role in determining the expression levels in the library. Secondary structure of mRNA is well-known to influence the rate of translation at initiation (16), elongation (31) and termination (32). In our library, because we randomized 12 nucleotides around the stop codon, variations in the secondary structure could potentially affect translation termination efficiency. However, no correlation was found between predicted folding energy around the stop codon and expression levels (Fig. S20).

In order to test if the impact of the C-terminal sequence was specific to the Dam protein, we measured the expression levels of a different reporter gene, (chloramphenicol acetyltransferase, *cat)*, when adding one of the 20 amino acids to the C-terminal (Fig. S19). The results were in qualitative agreement (Fig. S21, pearson correlation 0.58 weak promoter, 0.61 strong promoter), and the same general trends were found: hydrophobic amino acids led to lower expression than positively charged amino acids (Fig. S19 panel C, t-test independent, p < 0.015).

In conclusion, we showed experimentally that the identity of the last two amino acids impacts protein expression levels, with differences in protein abundance up to 4-fold, with lysine leading to the highest expression levels, and hydrophobic amino acids leading to the lowest levels.

## Discussion

Previous studies of biases in C-terminal composition considered a small number of bacterial species (5–7, 9, 10) which limited their level of resolution. In our study, we took advantage of the vast increase in the number of assembled bacterial genomes (20) to explore C-terminal composition biases across bacterial taxonomy. Here, we showed that the main pattern of composition biases at position -1, namely the enrichment for lysine and arginine, the depletion of threonine and the depletion of hydrophobic amino acids, is conserved across all bacterial phyla. In fact, the biases for lysine and against threonine were found in the vast majority of the 280 taxonomic clades at the family rank (Fig. S7). Noteworthily, we observed a strong anticorrelation across all phyla between Lys and Thr biases. Similar biases were also found in eukaryotes, where positively charged residues are enriched at the C-terminal, while threonine and glycine are depleted (8, 10, 33). Moreover, we showed that this bias pattern was found in all protein functional categories. Thus, our results suggest that the main C-terminal amino acid composition biases are conserved across bacterial species.

By analyzing C-terminal composition at a finer level of detail, we also revealed variations in biases depending on the stop codon context and the identity of the 3rd nucleotide. Intriguingly, we found that NNA codons were in general more favored, while NNG were more disfavoured in the context of an UGA stop codon. In bacteria, genes that overlap on the same polycistronic mRNA are common (34) (34% in *E. coli* MG1655 (35, 36)). Both genomic compression and translational coupling are believed to promote short overlaps of 4 nt, in which the use of the UGA stop codon is particularly frequent (37). Thus, the preference of the NNA codons in the UGA stop codon context could reflect selection constraints of overlapping start codons. Overall, biases were observed at multiple levels and are probably the result of multiple evolutionary constraints acting at the C-terminus.

We further found indications of a weak but significant selective pressure acting specifically on the identity of the C-terminal residue of highly abundant proteins. Both the enrichment for lysine and the depletion of threonine were stronger in highly abundant proteins, showing a clear association between protein abundance and C-terminal amino acid identity. In addition, hydrophobic amino acids Val, Ile, Leu and Met were disfavored in all proteins groups. In agreement with this, our expression assay revealed that C-terminal lysine, and to a lesser extent arginine, increased the expression levels of the *dam* reporter gene in *M. pneumoniae*, and that the presence of hydrophobic amino acids decreased expression levels. Qualitatively similar results were obtained with another reporter gene, supporting the notion that the observed modulations in expression are independent from the characteristics of the Dam protein. Altogether, our results suggest that the identity of the C-terminal amino acid modulates protein abundance in bacteria and is under selective pressure in highly abundant proteins. Which exact mechanism, or mechanisms, drive this phenomenon remains an open question. The fact that we see a gradient for the favoured and disfavoured amino acids, except for Thr and for Pro in some phyla, suggests at least two different mechanisms acting on the identity of the C-terminal residue.

Protein abundance level results from the balance between synthesis and degradation rates, and C-terminal sequence composition could potentially influence one or both of these processes.

One possible mechanism that could potentially drive the preferences for C-terminal amino acid and modulate protein abundance is protein degradation. In bacteria, apart from the universal N-terminal signals recognized by the N-end-rule pathway, several examples of degrons that target proteins to specific proteases have been identified (38). For instance, the *E. coli* ClpXP protease recognizes the ssrA tag at the C-terminal of proteins (13, 14). The ssrA tag sequence varies across bacterial species, but exhibit a similar C-terminal LAA-coo^-^ motif, with the exception of Mollicutes, where the ssrA tag sequences are unusually longer and exhibit a NYAFA-coo^-^ conserved C-terminal motif (39). *M. pneumoniae* does not possess ClpXP, but the ssrA tag is efficiently degraded *in vitro* by recombinant Lon protease (39). Interestingly, aspartic substitutions of the last two hydrophobic amino acids (FA-Stop) of the *M. pneumoniae* ssrA tag prevent the efficient degradation by recombinant Lon (40). Despite the importance of these C-terminal residues, several other amino acids within the ssrA tag seem to be also involved for efficient recognition and degradation, suggesting the existence of multiple signalling elements. In our analyses we found biases against hydrophobic amino acids at the last C-terminal position, which correlated with reduced levels of expression of two different gene reporters in *M. pneumoniae*. Interestingly, a similar observation was made for the halophilic archaeon *Haloferax volcanii (41)*, suggesting this could be a common mechanism to modulate protein levels. Whether this reduction in protein expression occurs post-translationally is unclear. However, protein degradation dependent on the universal Lon protease has been associated to degrons rich in aromatic and hydrophobic residues that would typically be buried in the hydrophobic core of native proteins (42).

Differential efficiency of translation termination is another possible mechanism that could affect protein synthesis rates. Proline at C-terminal can induce translation termination stalling in *E. coli* (11), which is usually alleviated by the *ssrA* system. Regarding threonine, one of the few C-terminal sequence motifs identified that induce termination stalling in *E. coli* (12) included Thr at position −1 with a hydrophobic-rich motif upstream (WILFXXT-Stop). Moreover, the threonine residue was shown to be essential for the stalling effect, suggesting that threonine might reduce termination efficiency in specific contexts. However, despite threonine was found to be depleted in all bacterial species and in the Mycoplasma genus (odds ratio 0.41), the measured expression level for variants with C-terminal threonine were close to the average. The same happened with proline (odds ratio 0.25 in *M.pneumoniae*).

Based on these observations, we postulate that the presence of threonine or proline at the C-terminal of a highly abundant protein could impact negatively the fitness of the bacterium without changing the protein expression level. For example, a moderate pausing at translation termination could increase the load of ribosomes on the mRNA while barely changing the total protein output, if termination is not the limiting rate in translation. The subsequent reduction in the pool of free ribosomes could decrease the expression of the other genes and lead to a fitness cost (16, 43). In this mechanism, the selection bias will be seen only at the −1 position and no gradient will be seen, which is the case for these amino acids.

The variability across taxonomy of the bias for proline also suggests that translation termination stalling at C-terminal proline could be species specific. In this regard, an *in vitro* study of the release factor (RF)-mediated rates of peptide release has suggested that peptide release is particularly inefficient on peptides terminating with proline or glycine, for both RF1 and RF2, and especially in the absence of methylated RFs factors (44). The addition of elongation factor P (EF-P) had no effect on the release rate suggesting that EF-P does not significantly promote peptide release on proline and glycine residues (44). On the other hand, prokaryotic release factor 3 (RF3) plays a role both in quality control during elongation (45) as well as in termination, where it promotes the recycling of RF1/2 and increases the rate of translation termination (46–49). While improving the efficiency and fidelity of termination, RF3 is not essential for growth in *E. coli* (50) and is absent from many groups of bacteria (51). Interestingly, we found that depletion of proline at C-terminal was more pronounced in taxonomic clades lacking RF3, like *M. pneumoniae* (t-test, p < 5.14e-4) (Fig. S27). This suggests that translation stalling at C-terminal proline might be dependent on the absence of RF3, which would explain the variability in proline biases across taxonomy.

In conclusion, our study reveals an important role of the C-terminal residues in protein expression. We propose that the preferences for C-terminal amino acid composition could be easily taken into account in the optimization of heterologous protein expression. Our results suggest that the addition or substitution of one or two lysine residues at the C-terminus of a protein sequence could increase expression, in particular when the original protein sequence contains hydrophobic amino acids at the C-terminus. Moreover, more subtle modulations in the protein expression levels, often needed in the design of synthetic circuits, could be achieved by controlling the identity of the C-terminal residues.

## Methods

### Bioinformatics analysis

#### Genomic and protein sequences

The main dataset in our analysis is the full set of annotated prokaryotic genomes of the NCBI RefSeq database (20, 52). We selected the RefSeq collection for several reasons: i) consistency and high quality in the gene annotation using a single annotation pipeline, ii) classification of species using the NCBI taxonomy, and iii) set of reference and representative genomes with curated annotation and balanced taxonomic diversity. We first downloaded the assembly summary file that reports metadata for the genome assemblies on the NCBI genomes FTP site at ftp://ftp.ncbi.nlm.nih.gov/genomes/refseq/assembly_summary_refseq.txt, on February 9, 2017. We selected the genomes in the representative or reference genome sets and with a complete genome assembly status (no contigs). All 1582 selected genome files were downloaded in Genbank format from the ftp website following ftp paths given in the summary file. For each genome, protein sequences together with codon sequences, stop codons and codon tables were extracted from the genome annotations.

#### Clustering

Protein families with highly similar sequences are very common in bacteria and are often the result of gene duplication. In our analysis, we seeked to avoid over-representation of duplicated proteins in the same bacterial species. In particular, duplicated proteins with identical C-terminal regions could bias the analysis of C-terminal amino acid composition. We applied a two-steps clustering to protein sequences in each bacterial species, to remove protein sequences that present both a high overall sequence identity and a high identity at their C-terminal region. We ran the CD-HIT clustering method (version 4.6) (53, 54) with 80% identity threshold of the overall protein sequences. The resulting clusters composed of more than one sequence were clustered again by running CD-HIT with 85% identity threshold on their C-terminal regions (20 last amino acid positions). We kept only one representative sequence for each of the final clusters. For each bacterial species, we thus obtain the set of non-redundant sequences. In addition, we also filter out proteins with length smaller than 50 residues. In total, we obtained 4 897 860 non-redundant sequences out of 4 934 952 (0.75% sequences removed).

#### Taxonomy

We used the ETE’s ncbi_taxonomy python module (55) which provides utilities to efficiently query the NCBI Taxonomy database (56). The NCBI Taxonomy database was downloaded as of February 2, 2017. For each genome in the assembly summary, the organism taxid was searched for in the taxonomic tree and taxonomic lineage information was retrieved. After removing non-bacterial genomes, we obtained 1582 genomes in the bacterial superkingdom, summing up 4 934 952 sequences.

Genomes were grouped by taxonomy at different ranks, ranging from superkingdom to species. In total, we obtained 2859 taxonomic groups which contained at least one genome. Our pipeline for C-terminal analysis was applied to the protein sequences sets of each of the taxonomic group, by grouping all non-redundant protein sequences in the taxon. We used the ETE package (55) to draw the taxonomic tree and the C-terminal biases of each clade.

#### Phylogenetic tree

We approximated evolutionary distances between the taxonomic phyla based on the bacterial phylogenetic tree of Lang et al (57), which used 761 bacterial taxa, inferred from a concatenated, partitioned alignment of 24 genes using RAxML. We mapped bacterial species to the NCBI taxonomy, obtaining 263 species with assigned phylum. While a phylogenetic tree will never be fully consistent with the taxonomy tree (58), we aimed at obtaining approximate evolutionary relationships at a high level of the taxonomy. Thus, we reconciled the NCBI phyla groups with the phylogenetic tree by pruning together monophyletic groups of species with the same phylum, discarded 38 species that were a minority in near-monophyletic groups. As recently described (58), the Tenericutes phylum should be included in the Firmicutes as a class. We relocated the Tenericutes phylum as a close sibling to Firmicutes. We obtained an approximated phylogenetic tree at the level of phyla.

#### Analysis of C-terminal bias

Amino acid composition of the C-terminal and N-terminal regions were analyzed as follows. Protein sequences were split into three parts: N-terminal region (20 first amino acids), C-terminal region (20 last amino acids) and bulk (remaining part of the sequence). The N-terminal region was discarded in order to remove well-known N-terminal sequence biases from the bulk composition. Position-dependent amino acid counts for the terminal regions were compared to bulk amino acid counts and two-tails Fisher exact test (scipy.stats package version 1.1.0 (59)) was used to test for the significance of enrichment or depletion of amino acid “X” at position *j* in the C-terminal region, where *j* ∈ [−20, …, −1]. The resulting odds ratios and p-values were computed for all 20 amino acids and 20 positions. Multiple tests correction was applied using the Benjamini-Hochberg method (from the statsmodels python package v0.9.0 (60)), with family-wise false discovery rate of 5%. Only significant biases after multiple tests correction were plotted. We followed the same method to study the bias in codon composition, and for the composition bias in amino acid pairs and codon pairs (hexamers) at the last two positions.

In the cases where the count of proteins was very low, even moderately strong biases would not appear as significant. Those cases of underpowered statistics were indicated in the plots with a hashed pattern. More precisely, based on the count of amino acid “X” in the bulk and its frequency, we computed the minimal count of expected observations of amino acid “X” at the C-terminal such that an odds ratio of 0.5 between observed and expected counts would lead to a p value lower than 0.05. For example, low frequency amino acids such as cysteine often had too low counts to be able to detect biases in the frequency at C-terminal.

In order to study the interaction between biases at two positions, or epistasis effect, we compared the frequency of an amino acid pair at the last two positions to its expected frequency under the assumption that both positions are independent (null model). More precisely, we defined the expected frequency as the product of the frequency of amino acid at position −2 and the frequency of amino acid at position −1. The deviation of the observed frequency from the expected was tested with the binomial test (scipy.stats.binom_test version 1.1.0).

#### Disordered regions

Prediction of disordered regions for 1305 bacterial genomes were downloaded from the D^2^P^2^ database (61) on 2019/07/31. Regions that were predicted as disordered by all six predictors (consensus of 100%) were considered.

#### Stop codon context

Protein sequences were classified by their stop codon context. Then, codon composition bias was analyzed as above within each class of stop codon context independently and multiple test corrections was applied within each class.

We also analyzed the relationship between the codon biases at position −1 and the identity of the codon third nucleotide in each stop codon context, by comparing the distribution of odds ratios of all codons for the selected 14 phyla (Fig. S10). We emphasize that the differences in biases that we found could not be explained by global variations in the 3rd nucleotide frequency, which could be influenced by the codon usage bias and genomic GC content of the bacterial species (62, 63), because each individual codon frequency at position −1 was compared to its frequency in the bulk.

#### Membrane proteins

We downloaded from the database PSORTb version 3.00 (64), which contains predicted subcellular localization for bacterial and archeal genomes, the full database tables for gram-positive and gram-negative bacteria. Based on RefSeq accession numbers, localization information could be assigned to 1 442 202 protein sequences out of 5 125 116 in our database. We then further selected 364 species in our database in which at least 80% of the proteins had localization information. Protein localized to the cytoplasmic membrane or to the outer membrane were grouped together as membrane proteins.

#### COG categories

In order to assign the Clusters of Orthologous Groups (COG) functional category to every protein in our NCBI-based database, eggnog-mapper version 0.12.7 (65) was run in the hidden Markov model (HMM) search mode against the bactNOG database. This mapper assigns functional annotation based on fast orthology assignments using precomputed clusters and phylogenies from the eggNOG database. The COG functional category is inferred from best matching orthologous group. The mapper failed to compute the assignment for 42 species. We could assign orthologous group to 4 213 953 protein sequences out of the total of 4 871 983 (86%).

#### Protein abundance

Protein abundance data sets were downloaded from the paxDB database release 4.1 (28). We selected the 24 bacterial species that were also present in our RefSeq-based database based on NCBI taxa id. Protein sequences were matched in two steps, first by matching paxDB ids to uniprot ids using the paxDB provided mapping file, then by matching uniprot ids to RefSeq ids using the uniprot mapping file. We obtained 62 093 RefSeq proteins (out of 78 483 in total) with at least one paxDB protein match. For the RefSeq proteins with multiple paxDB protein matches (1515 cases), the average protein abundance was computed. The resulting protein abundance coverage varied greatly among species. From the 24 species, only those where at least 40% of RefSeq protein sequences were assigned to an abundance value were kept (13 species, 40 002 RefSeq sequences, 26 935 with abundance value). Protein abundance values from the paxDB were given in relative values of part per million (ppm), which we assumed could be directly compared between species.

We then categorized proteins into three unequal bins: low abundance (0-20 percentiles), medium abundance (20-80 percentiles), and high abundance (80-100 percentiles). We applied the same analysis of bias as described above independently to the three groups of proteins. Note that the resulting biases at the C-terminal are relative to the amino acid bulk frequencies of each group of proteins, which may vary.

#### Conservation study

##### Orthogroup database

In order to define classes of highly and lowly abundant proteins, we followed a rough approximation by assuming that homologous proteins that are highly abundant in the 13 species with abundance data are highly abundant in all other species. More precisely, we selected orthogroups that had at least 10 protein sequences and at least 1 protein with abundance data, (13’388 groups), and classified them based on the abundance mean of all proteins within the group using the same low/medium/high abundance bins as above. We obtained 2084 orthogroups in the high abundance class, 7774 orthogroups in the medium abundance class, and 2454 orthogroups in the low abundance class.

##### Multiple sequence alignment

We analyzed the characteristics of the conservation profile of each orthogroup by computing their Multiple Sequence Alignment (MSA). For each orthogroup, redundant homologous sequences were first removed using the CD-HIT clustering method (53, 54) at 85% identity threshold. In order to limit the computation time for large orthogroups, 120 protein sequences were selected randomly for the MSA. MSAs were performed with MAFFT (66, 67), using the G-INS-i method (iterative refinement method with global alignment) and 1000 maximum iterations.

For some orthogroups, some sequences presented long extensions at the C-terminal. In order to allow for comparison within variable C-terminal regions of approximately similar lengths, we removed those sequences that presented an extended C-terminal domain longer than 25 residues, but only when those sequences summed up to less than 15% of the total number of sequences.

We then computed the conservation score of each MSA column as the Jensen-Shannon divergence (JSD) (68, 69), a popular measure of conservation. The JSD was computed in two ways: i) taking into account a gap penalty by multiplying the raw column JSD score by the fraction of non-gapped positions in the column (69, 70), and ii) removing gaps from the column (no gap penalty).

##### C-terminal variable region

The variable region was defined as the region of the MSA of low average conservation score at the C-terminal. For each orthogroup, the variable region was determined by finding the largest region starting from the C-terminal for which the average conservation score over a rolling window of 5 residues was always lower than a threshold of 0.3. Orthogroups with a non-conserved region of at least 6 residues long were then selected. In order to decrease the probability of the C-terminal being functional, we selected those regions with hyper-variable length. For each orthogroup, we computed the distribution of non-gapped residues sequence lengths within the C-terminal non-conserved region. The most variable region would present a uniform distribution, with an equal number of sequences for each possible length of its C-terminal region. The least variable region would be composed of all sequences having the same C-terminal region length. We computed the Kullback–Leibler divergence of the length distribution with respect to the uniform distribution, as a measure of the length variability of the region. Note that lengths with zero count do not contribute to the divergence, such that distributions with only few different lengths will have an artificially low divergence. We selected orthogroups that presented a divergence smaller than 1 and whose variable region presented at least 6 different lengths. We obtained 883 orthogroups with hyper-variable C-terminal region in the low abundance class (out of 2454) and 620 orthogroups in the high abundance class (out of 2069).

##### C-terminal-aligned conservation profile

For each orthogroup, we aligned sequences of the variable region to the C-terminal (right-aligned), and computed the Jensen-Shannon divergence (JSD) conservation score with no gap penalty at each column. In the absence of any selective pressure in the variable region, we would expect a low and constant conservation score at every position. However, due to our definition of the variable region, the gap fraction naturally decreases from more than 80% at the column −10 to approximately 10% at the last column (C-terminal). The JSD score with no gap penalty artificially increases when the number of non-gap residues in a column gets very small, such that only a few different amino acids are represented by chance. This is particularly true when the number of residues is below 40, as shown by the score distribution of randomly generated lists of amino acids of different lengths (Fig. S16). For most of the OGs, the number of non-gap residues in the right-aligned variable regions decreases rapidly below this threshold (Fig. S17). Thus, we expect that, in the absence of any selective pressure, the conservation score of right-aligned columns will constantly decrease from left to right.

In order to verify that the observed pattern was significant, we performed two controls. First, we shuffled the order of the residues within each variable region sequence, such that the exact amino acid composition within the variable sequence of each protein was maintained. We then recomputed the MSA, the position of the variable region and the C-terminal-aligned conservation profile. Second, we replaced each variable sequence with a random sequence of the same length, using the amino acid frequencies from the bulk of all sequences of all OGs in the corresponding class (high or low abundance). In both cases, the average conservation score at position −1 was lower than the one at position −2, in agreement with the null hypothesis. In the case of the random sequences, the decrease in JSD was highly significant (W.M.W. test p < 2.9e-3 and mean fold change 0.85 for low abundance class, W.M.W. test p < 9.0e-4 and mean fold change 0.81 for high abundance class). In the case of the shuffled sequences, although the mean decreased, the difference between the distributions at the two positions was not significant. This can be explained by the fact that for very short sequences, random shuffling did not always change the amino acid identity at the last two positions. For example, all variable sequences with a length of only one residue were not changed.

##### Amino acid frequency

First, we computed the bulk frequency of the OG class, by taking all the sequences in the OGs, removing the first 20 residues at the N-terminal and the C-terminal variable region. The a.a. frequency for each column in the C-terminal-aligned variable region was then normalized by the bulk frequency.

##### PDB structure annotation

Secondary structure blocks indicate alpha helix (red) and beta strand (gold) as predicted by STRIDE (71).

### Experimental assays

#### Construction of C-terminal *cat* derivatives and CAT expression quantification

Mini-transposon plasmids containing *cat* derivatives, in which one of the 20 amino acids were added to the C-terminal end were constructed as follows. The *cat* reporter gene was first amplified with primer cat_mp200Pr_F and the corresponding reverse primer listed in table S1. Each reverse primer incorporates at its 5’ end, the EcoRV restriction site and the codon sequence of one of the 20 amino acids. The corresponding PCR products were then used as templates for a second PCR, in which the cat_mp200Pr_F2 forward primer was used instead. This primer contains at its 5’ end the *Xho*I restriction site plus the mp200 promoter sequence to drive *cat* expression. Finally, all PCR products were cloned into a *Xho*I/*Eco*RV-digested pMTn*TetM438* mini-transposon vector (72).

To obtain *M. pneumoniae* M129 strains expressing the different C-terminal *cat* derivatives, all twenty constructs were transformed separately by electroporation as previously described (72) with few modifications. Briefly, 80 µl of cells (with approximately 10^9^ cells/ml) resuspended in electroporation buffer (8 mM HEPES·Na pH7.2, 272 mM sucrose) were mixed with 2 µg of DNA in a 1mm gapped electroporation cuvette (Bio-Rad). Cells were incubated on ice for 15 min and electroporated using the Gene Pulser XCell™ electroporation system (Bio-Rad) with the Pulse Controller set at 1.25 kV, 25 µF and 100 Ω. After 15 min of incubation on ice, cells were resuspended in 900 µl of Hayflick medium, incubated 2 h at 37°C, and the pool of transformants selected in Hayflick medium containing 2 μg/ml of tetracycline.

For quantification of CAT expression we used an ELISA based assay (CAT ELISA Kit assay, Roche). Briefly, *M. pneumoniae* transformant pools were grown at 37°C in 25 cm2 flasks containing 5 ml of Hayflick medium supplemented with 2 μg/ml of tetracycline. After 72 h, cells were washed three times in PBS, lysed with the CAT ELISA Kit lysis buffer and CAT expression determined following the manufacturer’s instructions. Approximately, 25 ng of cell lysate was used per ELISA plate well and absorbance recorded using a Tecan Infinite M200 plate reader. CAT expression was normalized by total protein amount, determined by the bicinchoninc acid (BCA) reagent Kit (Pierce).

#### ELM-seq

We followed the same protocol as described in Yus et al 2017 (19). Briefly, DNA adenine methylase (Dam) methylates multiple GATC sites that are located downstream of the *dam* coding sequence, in close proximity to the C-terminal randomized sequence. Dam expression level modulates the probability of having a site methylated, which is “measured” by differential digestion using methylation-sensitive restriction enzymes that cut the GATC sequence. Ultrasequencing of the region of interest allows to determine both sequence identity and expression level as reported by the ratio of reads between the two digested samples (DAMRatio).

#### Dam cloning and library preparation

A version of the minitransposon plasmid pMT85 was prepared by linker ligation, in order to accommodate the “reporter cassette” (i.e., the 4xGATC sites) at the C-terminus, followed by an endogenous (MPN517) terminator (so that all RNAs had a similar length). Briefly, two linker pairs, F_sp_term and R_sp_term, and F_term_NI and R_term_NI were annealed in two independent reactions, and ligated to pMT85 that had been open-cut with NsiI and EcoNI. In the next step the library was generated. The Gibson assembly reaction consisted in 3 pieces; the vector, cut-opened with NotI and Eco147I (introduced in a linker in the previous step), a PCR of dam, carrying the promoter in the forward oligos (F_P8_dam or F_mp200_dam and R_Cter_dam), and a linker with a C-terminus random region that was extended with Klenow in order to get the complementary random strand (F_damN_sp and R_damN_sp). The random region was ordered to have 40% GC content, similar to that of M. pneumoniae genome.

#### ELM-sequencing

Procedure for DNA-seq was as previously described (19), except for some details (Fig. S28). Contrary to the previous protocol, paired-end sequencing was chosen. In the final PCR step, a mix of equimolar forward oligos with a random region of increasing length (from 0-5 random nucleotides, see oligos: F_PE_Cdam, F_PE_n_Cdam, F_PE_n2_Cdam, F_PE_n3_Cdam, F_PE_n4_Cdam, F_PE_n5_Cdam), was designed in order to introduce enough diversity in the first Illumina DNA sequencing cycles. Two reverse oligos were used (R_PE_i6_Cdam and R_PE_i12_Cdam) alternatively, to be able to perform multiplexing. DpnI-treated DNA was amplified during 12 cycles, whereas MboI-treated DNA had to be amplified during 15 cycles.

#### DAMratio analysis

We followed the analysis pipeline described in (19). Raw reads were filtered for both the common C-terminal Dam-coding sequence and the varying downstream sequence according to the specific study as following CGCGAAAAAANNNNNNTAANNNNNNCAGGCCTTGA. In order to reduce the technical variability in the DAMRatio, we filtered out the sequences that did not have at least 30 reads in either one of the two enzyme-treated samples. In the end, 161227 and 194856 different sequence variants remained for strong and weak promoters, respectively. Higher thresholds resulted in the same or lower correlation between codon pair DAMRatios of the weak and strong promoter libraries.

Sequences that contained the GATC motif may introduce a bias in the measurement of the DAMratio and thus were filtered out prior to the analysis. Indeed, the number of GATC sites can modify the distribution of the DAMratios for a given library design (19). In the case of the weak promoter, sequences containing GATC showed a lower DAMratio than the other sequences. In the case of the strong promoter, the opposite trend was observed.

#### Epistasis

In order to compute the intensity of the cooperative effect between the identities of the last two amino acids on the expression level, we first assumed that the effect of each amino acid is independent, such that the expected expression level of an amino acid pair would be

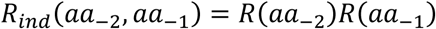

where *R*(*aa*_*i*_) is the relative expression level of all sequences with amino acid *aa* at position *i*, as measured by the DAMRatio normalized to the average. The cooperativity effect (or epistasis), was then computed as the ratio of the measured expression level of the pair to the expected one,

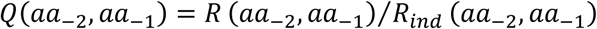

which in log scale reads,

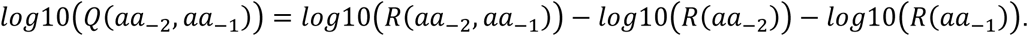

## Data and software availability

All the bioinformatics analysis were done using custom Python 3.6 scripts. Source codes are available upon request. Both the raw sequencing data and the processed data for the ELM-seq experiment are available at ArrayExpress with accession number E-MTAB-8223.

## Acknowledgments

We acknowledge the support of the Spanish Ministry of Economy, Industry and Competitiveness (MEIC) to the EMBL partnership, the Centro de Excelencia Severo Ochoa and the CERCA Programme / Generalitat de Catalunya. This project has received funding from the European Union’s Horizon 2020 research and innovation programme under grant agreement 634942 (MycoSynVac). ML-S acknowledges the support from FEDER project from Instituto Carlos III (ISCIII, Acción Estratégica en Salud 2016) (reference CP16/00094). The authors would like to thank the CRG Genomics Unit for assistance with the design and sequencing of the ELM-seq library.

## Author contributions

MW, RB, EY, JY, ML-S and LS conceived and designed the study. MW conceived and performed all bioinformatics analyses. EY designed the library and performed the ELM-seq experiment. MW and JY analyzed the ELM-seq results. RB designed and performed the Cat reporter assay. LS and ML-S provided direct supervision. MW, RB, JY, ML-S and LS wrote the manuscript. All authors read and approved the final manuscript.

